# Entorhinal-hippocampal circuit integrity is related to mnemonic discrimination and amyloid-β pathology in older adults

**DOI:** 10.1101/2022.05.10.491249

**Authors:** Jenna N. Adams, Soyun Kim, Batool Rizvi, Mithra Sathishkumar, Lisa Taylor, Alyssa L. Harris, Abanoub Mikhail, David B. Keator, Liv McMillan, Michael A. Yassa

## Abstract

Mnemonic discrimination, a cognitive process that relies on hippocampal pattern separation, is one of the first memory domains to decline in aging and preclinical Alzheimer’s disease. We tested if functional connectivity (FC) within the entorhinal-hippocampal circuit, measured with high-resolution resting state fMRI, is associated with mnemonic discrimination and Aβ pathology, measured with PET, in nondemented older adults. Low object mnemonic discrimination performance was specifically associated with increased FC between anterolateral entorhinal cortex (alEC) and dentate gyrus (DG)/CA3, supporting the importance of this connection to object memory. This hyperconnectivity between alEC-DG/CA3 was related to Aβ pathology and decreased entorhinal cortex volume. In contrast, spatial mnemonic discrimination was not associated with altered FC. Aβ was further associated with decreased FC and volume within hippocampal subfields. Our findings suggest that Aβ may indirectly lead to memory impairment through entorhinal-hippocampal circuit dysfunction and neurodegeneration, and provide a mechanism for vulnerability of object mnemonic discrimination.

## INTRODUCTION

Decline in episodic memory is a hallmark feature of both aging and Alzheimer’s disease (AD). Mnemonic discrimination, a process which supports episodic memory, has been shown to be particularly vulnerable to decline in older adults^1^. Mnemonic discrimination relies on pattern separation, the ability to differentiate similar memories into distinct, orthogonalized neural representations, which reduces interference and enables the encoding of highly similar experiences as unique episodes^2^. This neural computation requires intact processing in the dentate gyrus (DG) and CA3 subfield of the hippocampus and is dependent on direct input from the entorhinal cortex via the perforant pathway^2–5^. In aging, it is postulated that reduced inhibitory control in the DG and CA3 leads to

CA3 hyperactivation, driven by recurrent collaterals in this network^4^, resulting in a shift in computational balance away from pattern separation and towards pattern completion, or the retrieval of existing representations when prompted with partial or degraded cues^1, 6^. Previous studies have demonstrated that hyperactivation within DG/CA3 is associated with worse mnemonic discrimination performance in low-performing older adults and patients with amnestic mild cognitive impairment (aMCI)^7, 8^. Reduction of this hyperactivation with low doses of antiepileptic medication improves mnemonic discrimination performance^9^, suggesting that this increase in activity is an index of dysfunction.

Emerging research suggests that object and spatial mnemonic discrimination appear to be differentially vulnerable to aging. Performance on object mnemonic discrimination tasks, which test memory for changes in the object stimuli themselves, declines more sharply in older adults compared to performance on spatial mnemonic discrimination tasks, which test memory for changes in spatial position^10, 11^. The contributions of distinct information processing pathways within the entorhinal cortex, in which the anterolateral (alEC) and posteromedial (pmEC) subregions preferentially process object and spatial information, respectively, may explain this differential vulnerability^12–15^. Consistent with these behavioral findings, activity within the alEC is reduced in older compared to young adults during object mnemonic discrimination, with no change in pmEC activation during spatial mnemonic discrimination^16^.

One mechanism that may lead to dysfunctional information transfer between the entorhinal cortex and hippocampus in older adults is the development of amyloid-β (Aβ) and hyperphosphorylated tau, the pathological proteins characteristic of AD. The alEC is among the first cortical regions to develop tau pathology, which specifically targets the layer II cells that project to DG and CA3 as the perforant path^17, 18^. Aβ pathology has been shown to cause aberrant neural activity^19, 20^, including within the entorhinal cortex and hippocampus^21–23^, and promotes the development of tau in entorhinal cortex^24, 25^. Together, these two pathologies likely impede normal function of the entorhinal-hippocampal circuit, which may disrupt pattern separation mechanisms resulting in a decline in mnemonic discrimination, with the object domain particularly vulnerable to these effects due to the earlier susceptibility of alEC to the pathologies of Alzheimer’s disease.

Because Alzheimer’s pathology begins to develop while older adults are still cognitively normal^26^, it is critical to assess how the emergence of this pathology may lead to neural dysfunction in the medial temporal lobe and the initial expression of mnemonic discrimination deficits. Previous research in cognitively normal older adults has shown that Alzheimer’s pathology is related to both performance as well as activation while performing mnemonic discrimination tasks^27–29^. However, studies relating Alzheimer’s pathology to medial temporal lobe function during pattern separation have largely focused on measures reflecting the whole hippocampus, rather than the DG/CA3 region specifically, and task-based functional activation. Because communication between entorhinal cortex and DG/CA3 is critical to supporting pattern separation, assessing functional connectivity (FC) between subregions may provide more sensitive information about the integrity of this circuit, which is needed to fully understand decline in episodic memory in older adults.

The goal of the current study was to determine if impaired mnemonic discrimination performance in aging is related to altered FC within the entorhinal-hippocampal circuit, and whether altered FC may be attributed to the development of Alzheimer’s pathology. To test this, we conducted high-resolution resting state functional MRI on a sample of cognitively normal older adults who performed object and spatial versions of a mnemonic discrimination task outside the scanner. This allowed for a fine-grained examination of how FC between entorhinal subregions and hippocampal subfields is related to mnemonic discrimination performance. Due to distinct information processing pathways, we hypothesized that impaired object mnemonic discrimination would be related to altered FC between alEC and DG/CA3, while impaired spatial mnemonic discrimination would be associated with altered FC between pmEC and DG/CA3.

We then tested if Aβ positivity, an indication of future progression towards AD and likely concurrent tau pathology in entorhinal cortex^30^, was associated with FC within the entorhinal-hippocampal circuit. We hypothesized that Aβ positivity would be associated with altered FC between entorhinal cortex and DG/CA3, providing a potential mechanism for decline in mnemonic discrimination, as well as further FC disruption between hippocampal subfields due to its potential widespread impact on the circuit. Finally, to understand other factors that may lead to impairment in mnemonic discrimination, we tested whether neurodegeneration within the entorhinal-hippocampal circuit and FC within cortical memory networks were related to mnemonic discrimination and Aβ pathology.

## RESULTS

### Participants

We analyzed resting state fMRI data, structural MRI data, and 18F-florbetapir PET Aβ scans from a total of 64 cognitively normal older adults recruited from a community-based sample. Participants completed a comprehensive cognitive assessment battery including word list recall as well as the object and spatial mnemonic discrimination tasks. Full demographic information for the sample is included in **Table 1**.

**Table 1.** Demographic information and characteristics of the sample.

|  | N | M (SD) or n (%) |
| --- | --- | --- |
| Age (years) | 64 | 71.3 (6.4) |
| Sex (female) | 64 | 43 (67.2%) |
| Education (years) | 64 | 16.3 (2.3) |
| MMSE | 63 | 28.5 (1.3) |
| RAVLT Delayed Recall | 63 | 10.9 (3.3) |
| MDT Object LDI (high similarity) | 56 | 0.20 (0.13) |
| MDT Spatial LDI (high similarity) | 59 | 0.20 (0.14) |
| Aβ+ | 57 | 19 (33.3%) |
| Global FBP SUVR | 57 | 1.11 (0.19) |

All 64 participants received high resolution resting state fMRI, while subsets additionally received 18F-florbetapir-PET (FBP) and performed the mnemonic discrimination task (MDT). MMSE, Mini Mental State Exam; RAVLT, Rey Auditory Verbal Learning Task; LDI, lure discrimination index calculated on the high similarity stimuli; SUVR, standardized uptake value ratio.

### Mnemonic Discrimination Performance and Classification

Participants performed object (n=56) and spatial (n=59) versions of a mnemonic discrimination task (MDT; **Figure 1**). Full descriptions of the tasks are included in *Methods*. Briefly, participants first completed a study phase in which they were exposed to the target stimuli. In the test phase, participants judged whether stimuli were target or lure stimuli. In the object task (MDTO), lure stimuli consist of subtle changes to the object itself, while in the spatial task (MDTS), lure stimuli consist of changes in the location of the object on a grid. Each lure stimulus can be classified as low or high similarity to the original target stimulus. We focused on performance for the highly similar lure stimuli based on our prior work showing that pattern separation mechanisms in older adults are especially impaired for highly similar compared to less similar lures^31^. Performance was measured with the well-established lure discrimination index (LDI; p(“New or Different” | Lure) – p (“New or Different” | Target)), which corrects for potential response bias.

**Figure 1.**
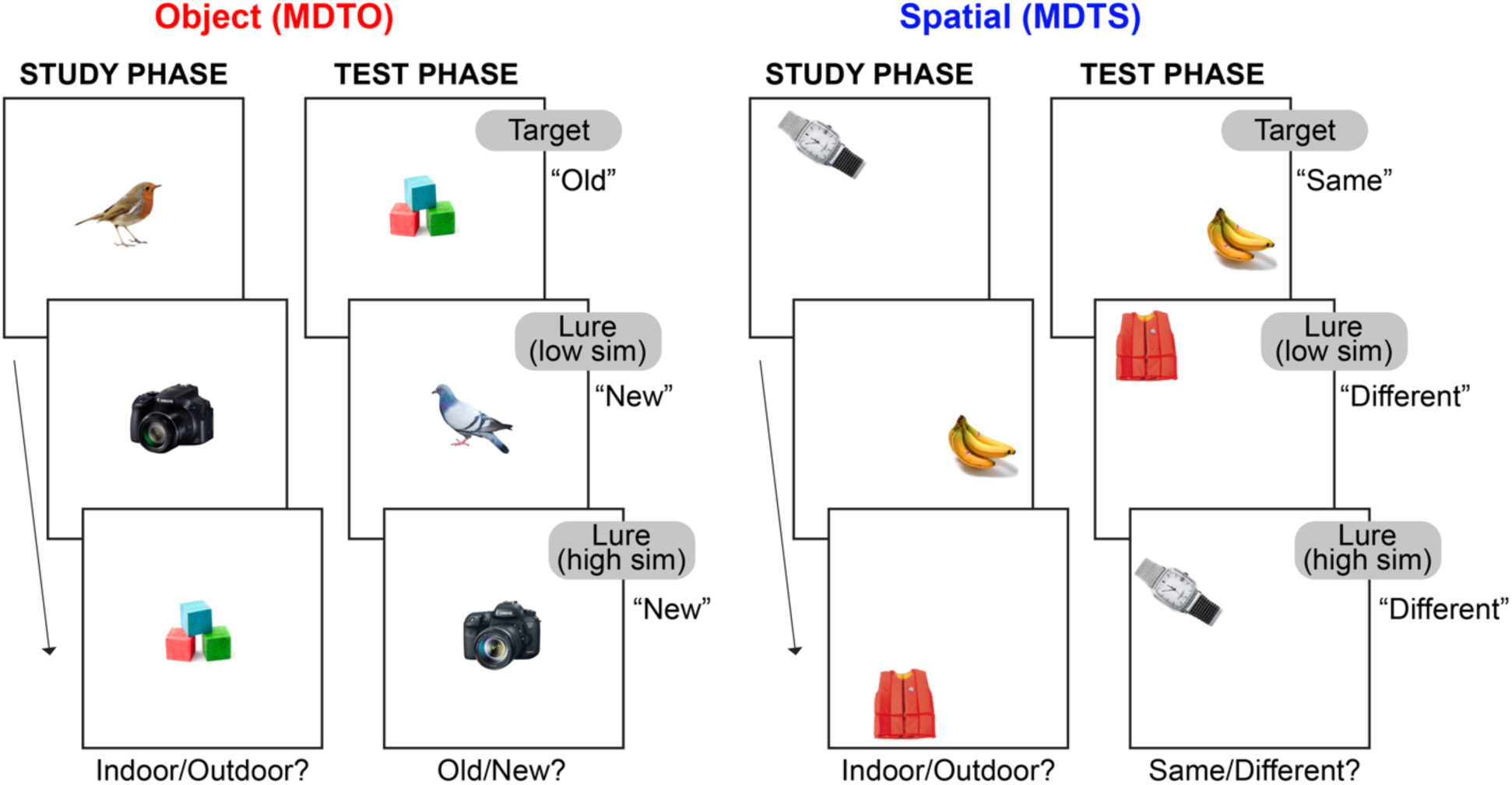
Object and spatial versions of the mnemonic discrimination task. Older adult participants completed both object (MDTO) and spatial (MDTS) versions of a mnemonic discrimination task. During the Study Phase, participants indicated whether each stimulus was more likely to be found indoors or outdoors. In the Test Phase, participants indicated whether each stimulus was “Old” (Target) or “New” (Lure) in the MDTO and “Same” (Target) or “Different” (Lure) in the MDTS. In the MDTO, lure stimuli consisted of variations to the object stimulus itself, while in the MDTS, lure stimuli were in a different spatial position on a grid. Lure stimuli in both tasks could be considered low similarity, where the differences were more readily detectable, or high similarity, where the changes were more subtle. We defined high and low performers on each task based on the lure discrimination index (p(“New or Different” | Lure) – p(“New or Different” | Target) computed from the highly similar lure stimuli, which maximally tax pattern separation mechanisms.

Performance on the object and spatial versions of the MDT were moderately correlated (r=0.46, p<0.001), and did not significantly differ (paired samples t-test, t(54)=0.34, p=0.73). There was no significant association between either object or spatial MDT with age or global FBP SUVR (ps>0.18), or sex difference in object MDT (p=0.53). However, female participants performed significantly higher than males on the spatial MDT (t(57)=2.13, p=0.04).

We classified participants as either low or high performers on the object and spatial MDT using a median split of the LDI high similarity score (MDTO, median=0.185; MDTS, median=0.21). For the object MDT, 28 participants (50%) were classified as high performers (MDTO+) and 28 participants (50%) classified as low performers (MDTO-). For the spatial MDT, 26 participants (44%) were classified as high performers (MDTS+) and 33 participants (56%) as low performers (MDTS-).

### Functional Connectivity

The primary aim of our study was to test how object and spatial mnemonic discrimination was related to FC between regions that comprise the canonical entorhinal-hippocampal circuit (entorhinal subregions and hippocampal subfields). We further investigated whether object and spatial mnemonic discrimination was related to FC between regions contained within the anterior-temporal and posterior-medial cortical networks^32^, respectively. This was achieved by analyzing high resolution (1.8mm^3^, partial brain acquisition) resting state fMRI data with a highly detailed atlas of regions of interest (ROIs) spanning the medial temporal lobe and hippocampus, shown in **Figure 2A**. We conducted ROI-to-ROI FC analyses using the CONN toolbox^33^. To ensure spatial specificity, BOLD time series were extracted from unsmoothed data. Further, we performed semipartial correlations between time series, which controls for signal from all other included ROIs, to mitigate any potential signal bleed in from neighboring regions and to obtain FC values representing the unique FC between regions^34, 35^.

**Figure 2.**
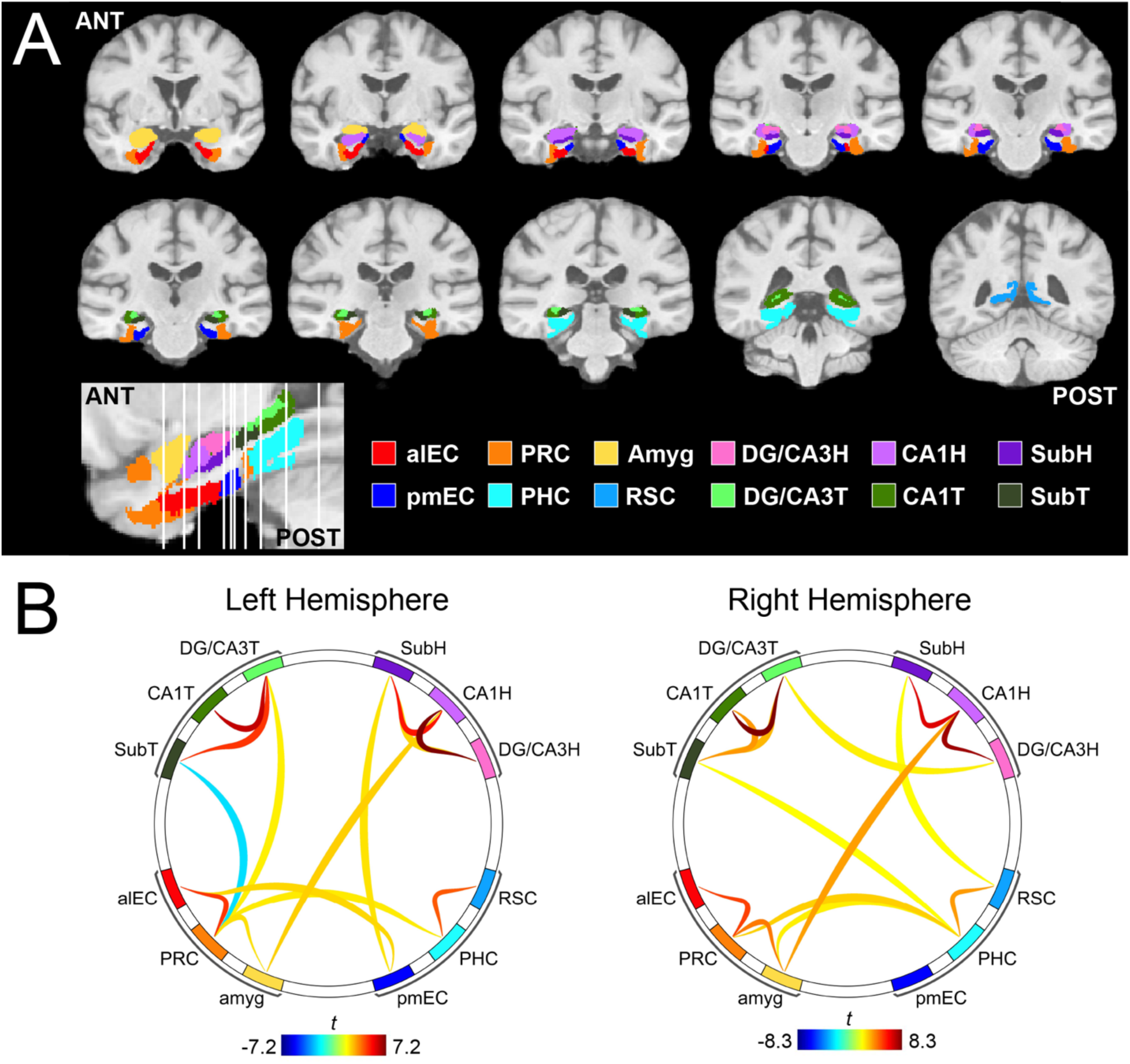
Functional connectivity (FC) methods and group-level results. We measured FC between regions of interest (ROIs) based on a high-resolution study-specific atlas which spanned the medial temporal lobes and related regions (**A**). ROIs included the anterior-temporal cortical network [anterolateral entorhinal cortex (alEC), perirhinal cortex (PRC), amygdala (amyg)], posterior-medial network [posteromedial entorhinal cortex (pmEC), parahippocampal gyrus (PHC), retrosplenial cortex (RSC)], as well as hippocampal subfields within the head (H) and tail (T) segments [dentate gyrus (DG)/CA3, CA1, and Subiculum (Sub)]. (**B**) Seed-to-seed FC was performed using semipartial correlations. Group-level results were thresholded using a connection threshold of p<0.05 p-uncorrected and ROI threshold of p<0.05 p-FDR corrected. FC was similar between hemispheres and predominantly occurred within networks.

Group-level patterns of FC are shown in **Figure 2B.** Overall, FC between the two hemispheres was largely consistent, and was strongest within each network (hippocampus, anterior-temporal, and posterior-medial networks). Within the hippocampus, there was strong FC between DG/CA3 and CA1, and between CA1 and subiculum (Sub), mirroring the known anatomical connectivity of this circuit. Because we did not have hypotheses regarding lateralized hemispheric effects, and to reduce the number of statistical tests, we averaged FC values between left and right hemisphere region pairs prior to subsequent analyses.

### Impaired object mnemonic discrimination is related to alEC-DG/CA3 hyperconnectivity

We first tested whether object and spatial mnemonic discrimination were associated with FC within the entorhinal-hippocampal circuit. We focused on FC between region pairs that compose the canonical entorhinal-hippocampal circuit: alEC to DG/CA3, pmEC to DG/CA3, DG/CA3 to CA1, and CA1 to Sub. While we acknowledge that FC is a non-directional measure, projections within the trisynaptic circuit are largely unidirectional^3,36^ and are likely the primary drivers of the FC measure. Primary analyses focused on subfields within the hippocampal head due to the increased vulnerability of this segment to aging and AD^37, 38^. We conducted two repeated measures ANCOVAs, one for each task, with pairwise FC measures (alEC-DG/CA3H; pmEC-DG/CA3H; DG/CA3H-CA1H; CA1H-SubH) within subjects and group assignment (high vs. low performers) between subjects, and age and sex as covariates of no interest. Our effects of interest were the main effect of group, indicating that FC differed between the groups uniformly across the circuit, and a group by FC interaction, indicating that FC differed between the groups in specific connections.

We first compared entorhinal-hippocampal circuit FC between low and high performers on the object mnemonic discrimination task (**Figure 3A**). While there was no main effect of performance (F(1)=2.27, p=0.14), we found a significant performance by FC interaction (F(3)=3.35, p=0.02), indicating the groups differed in FC in particular connections within the circuit. Post-hoc pairwise comparisons indicated that this interaction was driven by low performers having significantly increased FC between alEC and DG/CA3H compared to high performers (t(52)=2.81, p=0.007). There were no other significant group differences across the circuit (ps>0.30). The specificity of this result supports our hypothesis that the connection between alEC and DG/CA3H supports object mnemonic discrimination and its integrity tracks with performance.

**Figure 3.**
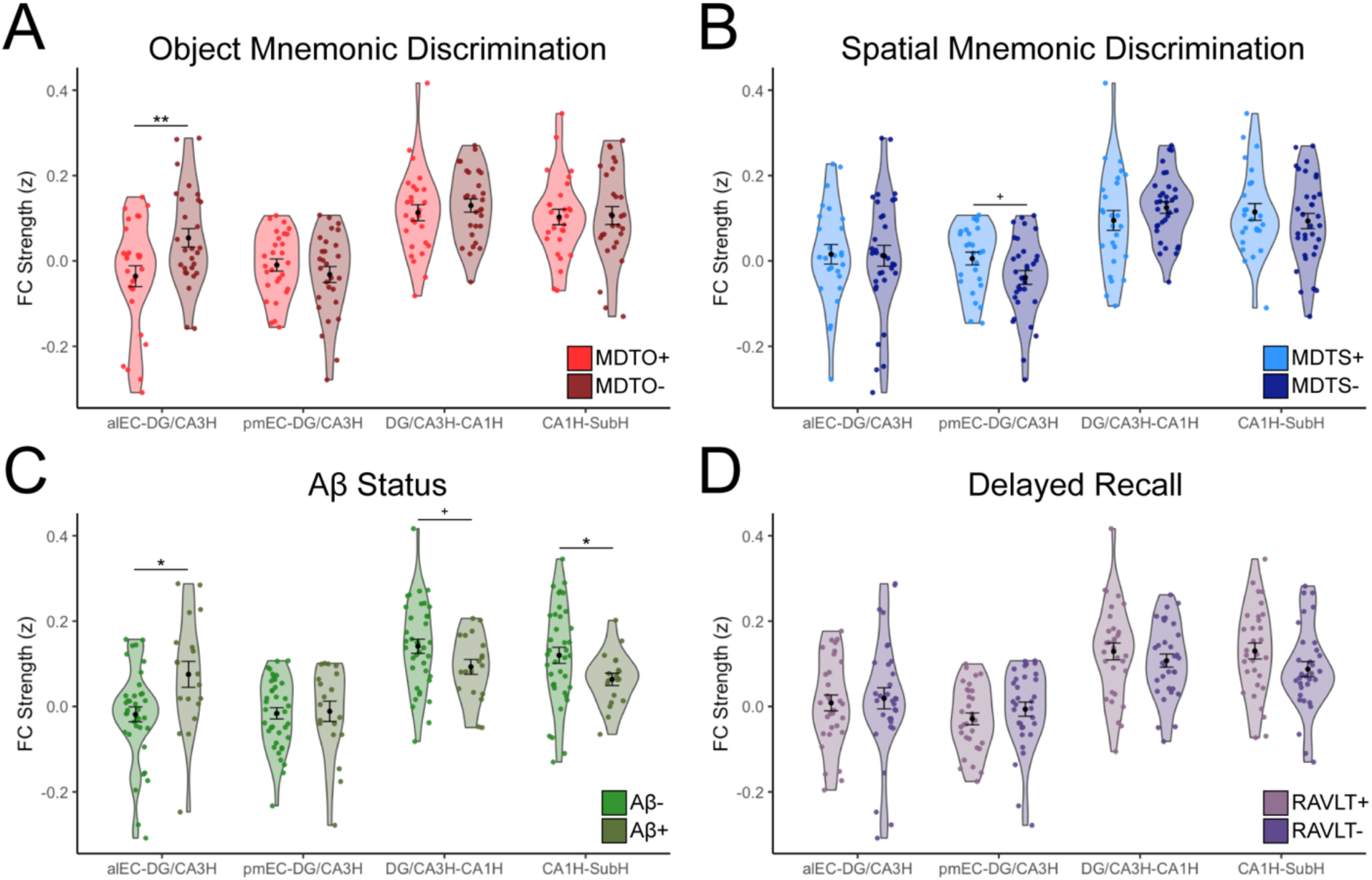
Entorhinal-hippocampal functional connectivity (FC) is related to object mnemonic discrimination and Aβ status. FC strength (Fisher z-transformed semi-partial correlation values) was calculated between each region pair corresponding to canonical anatomical connections within entorhinal-hippocampal circuit. (**A**) FC strength was compared between high (MDTO+; light red) and low (MDTO-; dark red) performers on the object mnemonic discrimination task (MDTO). Low performers had significantly increased FC between alEC-DG/CA3H compared to high performers, but no other differences across the circuit. (**B**) FC strength was compared between high (MDTS+; light blue) and low (MDTS-; dark blue) performers on the spatial mnemonic discrimination task (MDTS). There were no significant differences in FC strength, however, there was a trend-level association for decreased pmEC-DG/CA3H FC in the low compared to high performers. (**C**) FC strength was compared between Aβ- (light green) and Aβ+ (dark green) participants. Aβ+ participants had significantly increased FC between alEC-DG/CA3H compared to Aβ- participants, consistent with the pattern found in the MDTO- group. Aβ+ participants also had decreased FC between CA1H-SubH, and a trend-level association for decreased FC between DG/CA3H-CA1H compared Aβ- participants. (**D**) Control analyses tested whether performance on a traditional episodic memory measure, the Rey Auditory Learning Verbal Test (RAVLT) delayed recall, was associated with changes in entorhinal-hippocampal FC. There was no difference in FC between high (RAVLT+; light purple) and low (RAVLT-; dark purple) performers, indicating specific effects to mnemonic discrimination. alEC, anterolateral entorhinal cortex; pmEC, posteromedial entorhinal cortex; DG/CA3H, dentate gyrus/CA3 head; CA1H, CA1 head; SubH, subiculum head. Error bars represent standard error of the mean. **p<0.01 *p<0.05 ^+^p<0.10

We next compared entorhinal-hippocampal FC between low and high performers on the spatial mnemonic discrimination task (**Figure 3B**). We did not observe a significant main effect of performance (F(1)=0.81, p=0.37) or performance by FC interaction (F(3)=1.01, p=0.39), indicating that there were no general or regionally specific differences in entorhinal-hippocampal circuit FC related to spatial mnemonic discrimination performance. However, a targeted analysis of the connection between pmEC-DG/CA3H, which we specifically hypothesized would be related to spatial mnemonic discrimination, identified a trend in which the low performers had decreased FC compared to high performers (t(55)=-2.0, p=0.051).

We then examined whether these effects generalized to the hippocampal tail. We conducted similar ANCOVA models, instead including FC between entorhinal subregions and hippocampal tail ROIs (alEC-DG/CA3T, pmEC-DG/CA3T, DG/CA3T-CA1T, CA1T-SubT). We found no significant main effect or interaction for the hippocampal tail for either object (main effect of performance: F(1)=0.72, p=0.40; performance by FC interaction: F(3)=1.38, p=0.25) or spatial (main effect of performance: F(1)=1.20, p=0.28; performance by FC interaction: F(3)=0.37, p=0.78) mnemonic discrimination performance, suggesting the increase in FC between alEC and DG/CA3 in low object mnemonic discrimination group was specific to the hippocampal head.

As a control analysis, we examined whether a general episodic memory performance, rather than mnemonic discrimination, would also be related to differences in alEC-DG/CA3H FC or any other connection within the circuit (**Figure 3D**). To test this, we classified participants as low or high performers on the RAVLT delayed recall test using a median split and performed a similar ANCOVA analysis. There was no significant main effect of performance (F(1)=0.46, p=0.50) or performance by FC interaction (F(3)=1.49, p=0.22) when splitting the groups by delayed recall episodic memory performance, supporting the specificity of alEC-DG/CA3H connectivity in supporting mnemonic discrimination.

Finally, we examined whether splitting MDT performance based on all lure similarity trial types, rather than focusing on highly similar lure stimuli, would produce consistent results. For object mnemonic discrimination, we still observed a significant performance by FC interaction (F(3)=3.39, p=0.02), again driven by increased FC between alEC and DG/CA3H in low compared to high performers (t(52)=2.07, p=0.04), however this difference was less pronounced than when defining low performance only using high similarity lure stimuli. Again there were no significant effects when splitting spatial performance using all lure similarity trials (main effect of performance F(1)=0.005, p=0.95; performance by FC interaction F(3)=1.27, p=0.29).

### Aβ is associated with widespread changes in entorhinal-hippocampal circuit FC

Our second study goal was to determine whether Alzheimer’s pathology was also associated with changes to entorhinal-hippocampal circuit FC as a potential mechanism leading to mnemonic discrimination impairment. To assess this, we classified participants as Aβ- (n=38) or Aβ+ (n=19) using a threshold of global FBP SUVR >1.11^39^. While we did not have a direct measure of tau pathology, older adults classified as Aβ+ are highly likely to have tau pathology within the entorhinal cortex^30^. We then conducted repeated measures ANCOVAs, including FC (alEC-DG/CA3H; pmEC-DG/CA3H; DG/CA3H-CA1H; CA1H-SubH) as a repeated within-subjects measure, Aβ status (positive vs. negative) as a between-subjects factor, and age and sex as covariates of no interest, again with either a main effect of group or group by FC interaction as the effects of interest.

While there was no main effect of Aβ status (F(1)=0.05, p=0.82), we found a significant Aβ status by FC interaction (F(3)=5.73, p<0.001), indicating Aβ- and Aβ+ groups differed in FC in particular connections within the circuit (**Figure 3C**). Post-hoc pairwise comparisons indicated that this interaction was driven by widespread differences across the circuit. Aβ+ participants had significantly increased FC between alEC and DG/CA3H compared to Aβ- participants (t(53)=-2.67, p=0.01). This increase in alEC-DG/CA3H FC in the Aβ+ group mirrors the hyperconnectivity found in the low object mnemonic discrimination group. Further, Aβ+ participants had decreased FC between hippocampal subfields. Aβ+ participants had significantly decreased FC between CA1H and SubH compared to the Aβ- participants (t(53)=2.21, p=0.03), and a trend towards decreased FC between DG/CA3H and CA1H (t(53)=1.71, p=0.09).

We next tested the effect of Aβ on FC between the entorhinal subregions and subfields within the hippocampal tail. While there was no significant main effect of Aβ status (F(1)=0.07, p=0.80), there was a trend for a Aβ status by FC interaction (F(3)=2.48, p=0.06); however, because this effect did not reach statistical significance, we did not conduct further post-hoc pairwise comparisons.

### Contributions of anterior-temporal and posterior-medial cortical networks

Because the anterior-temporal network preferentially supports object processing, while the posterior-medial network preferentially supports spatial processing^32^, we investigated whether FC within these networks was related to object and spatial mnemonic discrimination, respectively. We included alEC, perirhinal cortex (PRC), and amygdala (amyg) to represent the anterior-temporal network, and pmEC, parahippocampal gyrus (PHG), and retrosplenial cortex (RSC) to represent the posterior-medial network. We acknowledge that additional regions are commonly included within these networks^32^, however, we chose to only include ROIs contained within our study specific template and were somewhat restricted in cortical coverage given our partial field of view in high-resolution scans.

To investigate the relationship between object mnemonic discrimination and the anterior-temporal network, we conducted a repeated-measures ANCOVA with FC (alEC- PRC, alEC-amyg, PRC-amyg) as a repeated within-subjects measure, performance (high vs. low performers) as a between-subjects factor, and age and sex as covariates of no interest. There was no significant main effect of performance (F(1) = 0.33, p = 0.57) or performance by FC interaction (F(2)=0.77, p=0.46), indicating connectivity between these regions did not strongly contribute to object mnemonic discrimination performance. We conducted a similar repeated-measures ANCOVA for spatial mnemonic discrimination and the posterior-medial network (pmEC-PHG, pmEC-RSC, PHG-RSC). Again, there was no main effect of performance (F(1)=0.38, p=0.54) or performance by FC interaction (F(2)=0.21, p=0.81).

Finally, we investigated whether Aβ positivity was associated with changes in FC within either network, based on evidence that Aβ accumulates in the posterior-medial network^29^ and can drive differences in connectivity within these networks^40, 41^. In the posterior-medial network, while there was no main effect of Aβ status (F(1)=0.36, p=0.55), there was a significant Aβ status by FC interaction (F(2)=3.99, p=0.02), indicating that FC within this network varied by Aβ status. Post-hoc pairwise comparisons indicated that this interaction was driven by the Aβ+ group having lower FC between pmEC and PHC (t(53)=2.49, p=0.02). Within the anterior-temporal cortical network, there was no significant main effect of Aβ status (F(1)=2.81, p=0.10) or Aβ status by FC interaction (F(2)=0.22, p=0.80).

### Factors associated with alEC-DG/CA3H hyperconnectivity

Due to the strong association of increased FC between alEC and DG/CA3 with both low object mnemonic discrimination performance and Aβ positivity, we conducted follow-up analyses to further explore factors related to this hyperconnectivity. We first examined whether alEC-DG/CA3H FC was continuously related to both performance on object mnemonic discrimination and levels of Aβ (global FBP SUVR). While the continuous association between object mnemonic discrimination and alEC-DG/CA3H FC did not reach significance (r=-0.19, p=0.17), we found a strong continuous association between higher global FBP SUVR and increased alEC-DG/CA3H FC (r=0.35, p=0.008; **Figure 4A**), which remained significant when controlling for age and sex (r=0.34, p=0.01).

**Figure 4.**
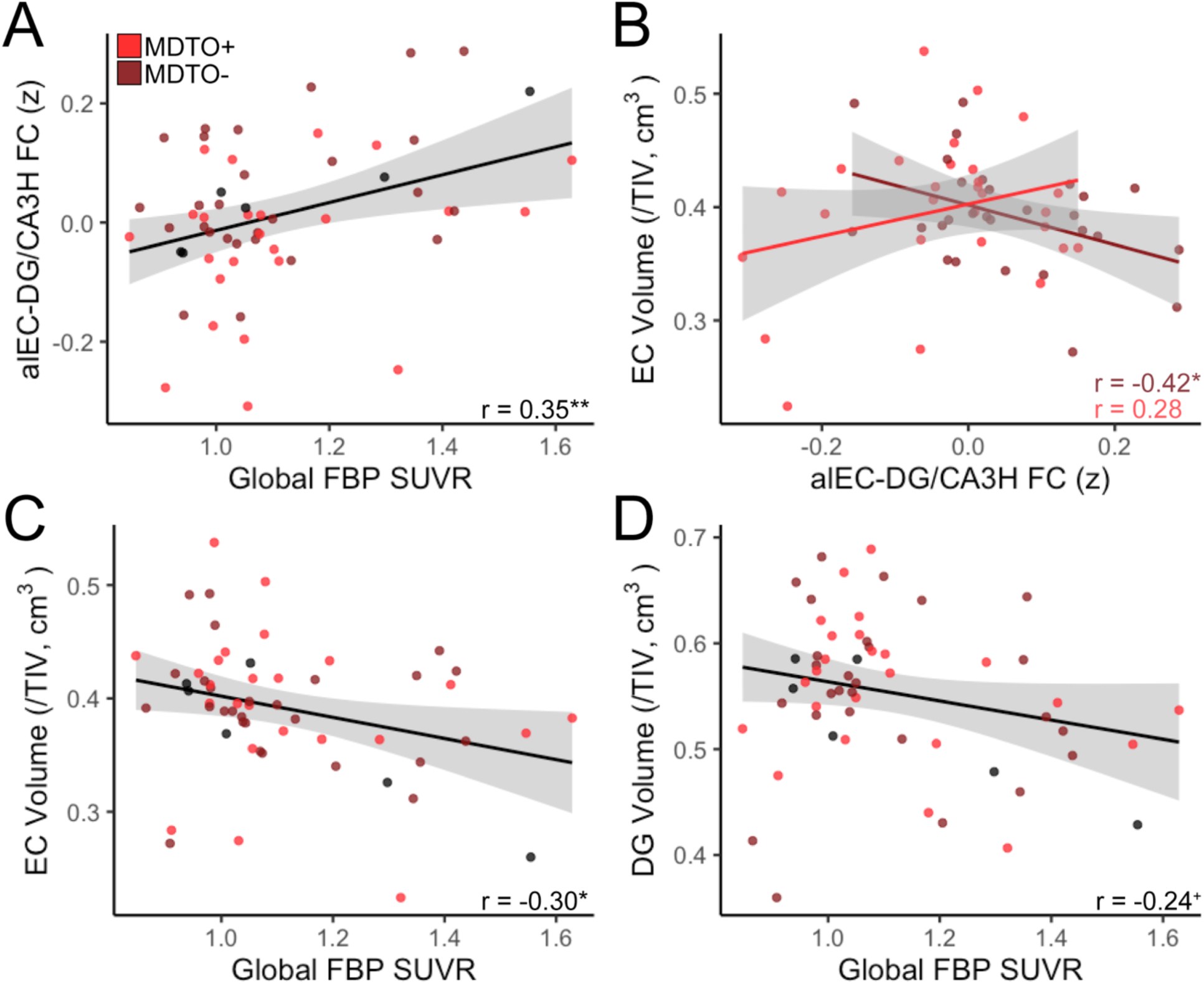
Associations between alEC-DG/CA3H hyperconnectivity with Aβ and neurodegeneration. (**A**) Increased alEC-DG/CA3H FC was significantly associated with continuous levels of global Aβ pathology (FBP SUVR). (**B**) There was a significant interaction between object mnemonic discrimination performance and neurodegeneration in predicting alEC-DG/CA3H FC. In the low object mnemonic discrimination group (MDTO-; light red), increased FC between alEC-DG/CA3H was associated with decreased EC volume, indicating neurodegeneration. There was no significant relationship in the high performing group (MDTO+; dark red). (**C**) Increased global FBP SUVR, indicating higher levels of Aβ pathology, was associated with reduced EC volume. (**D**) Increased global FBP SUVR had a trend-level association with reduced DG volume. alEC, anterolateral entorhinal cortex; DG/CA3H, dentate gyrus/CA3 head; EC, entorhinal cortex; DG, dentate gyrus; FBP, Florbetapir; TIV, total intracranial volume. **p<0.01 *p<0.05 ^+^p<0.10

We next investigated if alEC-DG/CA3H hyperconnectivity was related to neurodegeneration in these regions. To test this, we calculated native-space volume estimates using ASHS software, corrected for total intracranial volume (see ***Methods***). Because ASHS segmentation does not provide separate estimates for alEC and pmEC or for hippocampal head versus tail, we used the total volume of EC, DG, and CA3 in our analyses. Across all participants, there was no significant relationship between alEC- DG/CA3H FC and the volume of these regions (EC: r =-0.09, p=0.49; DG: r=-0.14, p=0.29; CA3: r=-0.16, p=0.21). However, we found a significant interaction between object mnemonic discrimination performance and EC volume in predicting alEC-DG/CA3H FC (model: F(50)=3.04, p=0.02; interaction t=2.51, p=0.02; **Figure 4B**). Investigating this interaction further, we found that within the low object mnemonic discrimination group, increased alEC-DG/CA3H FC was related to decreased EC volume (r=-0.42, p=0.03; controlling for age and sex: r=-0.38, p=0.06), while there was no significant relationship in the high object mnemonic discrimination group (r=0.28, p=0.15). This interaction was specific to EC volume; we did not observe a similar effect with DG or CA3 volume (model ps>0.10; interaction ps>0.30). Finally, performance on the object mnemonic discrimination task (both continuous associations and group comparisons) was not directly associated with volume in EC, DG, or CA3 (ps>0.55).

Because Aβ pathology was also related to alEC-DG/CA3 hyperconnectivity, we next tested whether Aβ was associated with neurodegeneration in these regions. Comparing volume of these regions in Aβ+ and Aβ- groups indicated that Aβ+ older adults had reduced volume in both EC (t(55)=2.44, p=0.02) and DG (t(55)=2.97, p=0.004), but not CA3 (t(55)=0.58, p=0.56). Further, a higher level of global FBP SUVR was associated with decreased volume in EC (r=-0.30, p=0.03; controlling for age and sex: r=-0.27, p=0.04; **Figure 4C**), with a trend for decreased volume in DG (r=-0.24, p=0.08; controlling for age and sex: r=-0.22, p=0.11; **Figure 4D**) and no significant relationship in CA3 (r=- 0.08, p=0.56). Together, these results indicate that the volume of the entorhinal cortex is closely associated with increased FC between alEC-DG/CA3H in low object mnemonic discrimination performers as well as Aβ pathology.

### Factors associated with CA1H-SubH hypoconnectivity

Because FC between CA1H and SubH was found to be decreased in Aβ+ older adults, we also explored how levels of Aβ and neurodegeneration were related to this hypoconnectivity. We found trend-level associations between decreased CA1H-SubH FC and both higher global FBP SUVR (r=-0.25, p=0.06; controlling for age and sex: r=-0.26, p=0.052; **Figure 5A**) and decreased CA1 volume (r=0.22, p=0.09; controlling for age and sex: r=0.22, p=0.08; **Figure 5B**), but no relationship with Sub volume (r=0.09, p=0.50; controlling for age and sex: r=0.08, p=0.52).

**Figure 5.**
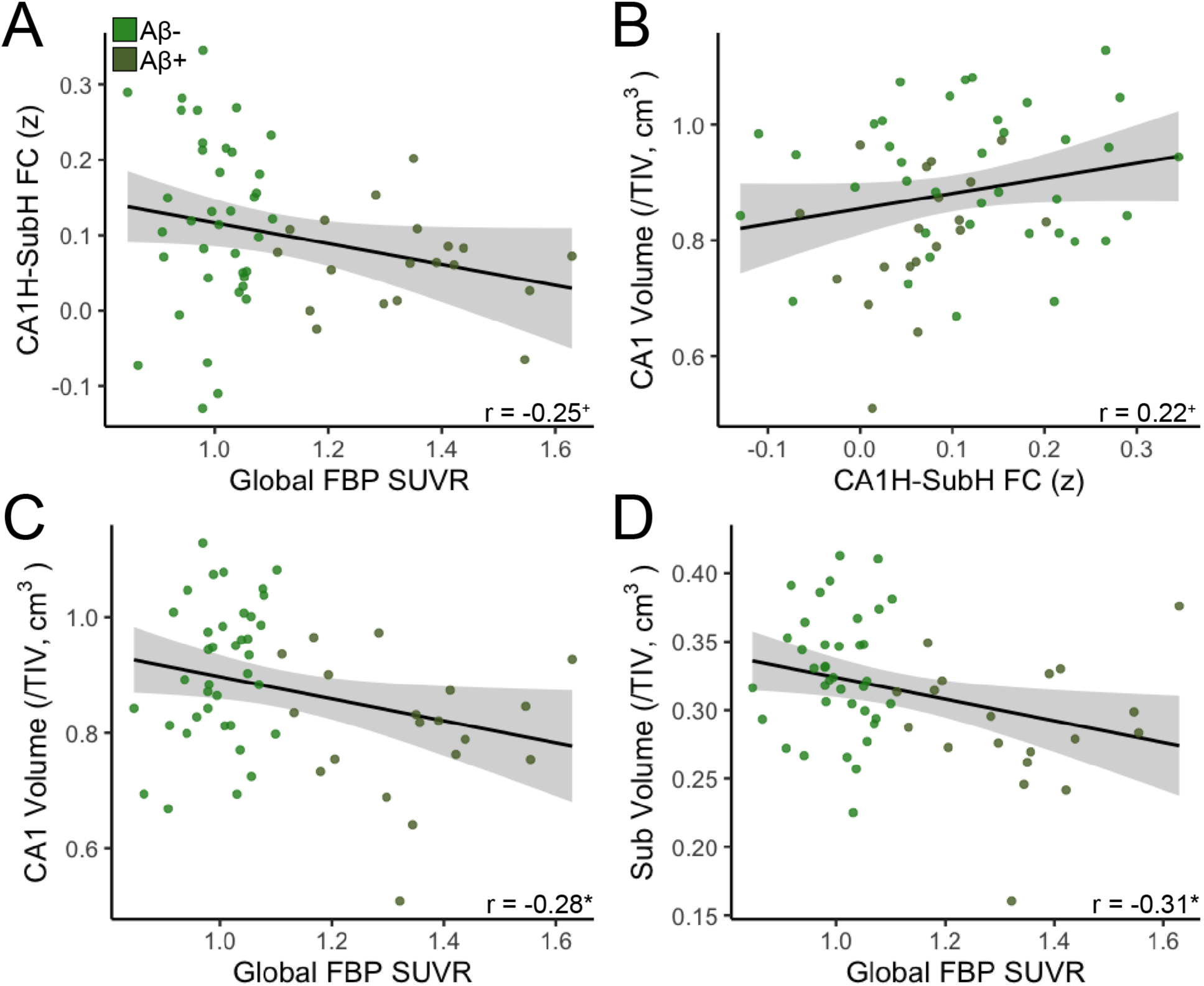
Associations between CA1H-SubH hypoconnectivity with Aβ and neurodegeneration. (**A**) There was a trend-level association between decreased CA1H-SubH FC with increased levels of global Aβ pathology (FBP SUVR). (**B**) There was a trend-level association between decreased CA1H- SubH FC and CA1 volume. (**C**) Increased FBP SUVR was significantly associated with reduced CA1 volume. (**D**) Increased FBP SUVR was significantly associated with reduced Sub volume. CA1H, CA1 head; SubH, subiculum head; Sub, subiculum; FBP, florbetapir; TIV, total intracranial volume. *p<0.05 ^+^p<0.10

Next, we investigated relationships between Aβ pathology and neurodegeneration in CA1 and Sub. Aβ+ older adults had reduced volume in CA1 (t(55)=3.05, p=0.004) and Sub (t(55)=3.02, p=0.004) compared to Aβ- older adults. Further, higher global FBP SUVR was continuously associated with decreased volume in CA1 (r=-0.28, p=0.03; controlling for age and sex: r=-0.29, p=0.03; **Figure 5C**) and Sub (r=-0.31, p=0.02; controlling for age and sex: r=-0.34, p=0.01; **Figure 5D**). These results suggest strong relationships between Aβ pathology and volume of both CA1 and Sub.

## DISCUSSION

Our results provide evidence that the integrity of communication within the entorhinal-hippocampal circuit, measured with resting state FC, is related to mnemonic discrimination performance and Aβ pathology in older adults who are cognitively normal. We found that hyperconnectivity between alEC and DG/CA3, particularly within the hippocampal head, is related to both poor object mnemonic discrimination performance and Aβ pathology. Increased FC in the low object mnemonic discrimination performers was also associated with neurodegeneration within entorhinal cortex, but not DG or CA3, suggesting entorhinal dysfunction may primarily drive hyperconnectivity. There were no significant FC differences underlying spatial mnemonic discrimination, suggesting spatial memory deficits may arise later in disease progression. Further, Aβ pathology was associated with hypoconnectivity between CA1 and Sub, as well as neurodegeneration within these regions, suggesting that Aβ has widespread effects on the integrity of the entorhinal-hippocampal circuit. Together, our findings indicate that specific alterations in communication between alEC and DG/CA3 contribute to the vulnerability of object mnemonic discrimination in aging, which may be attributable to underlying Alzheimer’s pathology.

To our knowledge, our study is the first examination of high-resolution resting state FC between entorhinal subregions and hippocampal subfields with mnemonic discrimination in a sample of cognitively normal older adults. We demonstrate that mnemonic discrimination performance for object stimuli was specifically related to hyperconnectivity between alEC and DG/CA3, particularly within the hippocampal head. This finding supports both human and animal data showing that the perforant path connecting entorhinal cortex and dentate gyrus is essential to mnemonic discrimination^31, 42, 43^. However, we extend previous findings by showing that object mnemonic discrimination performance is localized to connectivity between DG/CA3 and alEC, rather than pmEC, which is consistent with models of distinct processing streams in entorhinal cortex^14^. The specificity of the connection between alEC and DG/CA3 for mnemonic discrimination is further supported by our control analysis showing that this connection was not associated with performance on a delayed recall measure of episodic memory. While input from the entorhinal cortex to the hippocampus is critical for all hippocampal function^3^, subtle changes in this connection may first emerge as deficits in mnemonic discrimination that can be observed while older adults are still cognitively normal. This further supports the notion that mnemonic discrimination performance is a highly sensitive marker of emerging cognitive decline.

Our findings extend previous task-based fMRI findings that investigated activation in entorhinal subregions and hippocampal subfields during mnemonic discrimination in aging^16, 28, 44, 45^. In previous work, impaired object mnemonic discrimination in older adults was associated with relative hypoactivity within alEC and hyperactivity within DG/CA3 compared to young control participants^16^. The amount of hypoactivation in alEC was correlated with hyperactivation in DG/CA3, suggesting a functional link between the two processes. We interpret our finding of increased FC between alEC and DG/CA3 to reflect coordinated hyperactivation in the entorhinal-hippocampal circuit at rest, while alEC hypoactivation may emerge during active mnemonic discrimination processing when analyzing specific contrasts or group comparisons^16, 45^.

Because increased FC was associated with Aβ pathology and neurodegeneration, it suggests that hyperconnectivity is dysfunctional rather than compensatory. A pathological increase in FC between the entorhinal cortex and hippocampus at rest is supported by previous findings^38, 40, 46, 47^. For example, studies comparing resting state FC between patients with MCI or AD with elderly controls have found increased FC between entorhinal cortex and the hippocampus, particularly with the hippocampal head^38^, consistent with our results. Another study found increased FC between the entorhinal cortex and hippocampal subfields in healthy older African Americans at genetic risk for AD, which was further associated with impaired ability to generalize previously learned information^46^.

Our study provides evidence that hyperconnectivity between the entorhinal cortex and hippocampus is related to Aβ pathology. Findings from animal models suggest Aβ increases neural synchrony and elicits epileptiform activity^19, 20^, which may express as increased FC with resting state fMRI. Further, research suggests that Aβ may particularly affect GABAergic inhibition^20^, which may lead to dysfunction of inhibitory mechanisms within DG and lead to hyperactivation. Supporting these findings from animal models, in human neuroimaging studies Aβ pathology is associated with hyperactivation in both the hippocampus and entorhinal cortex^21–23^ and hyperconnectivity between cortical networks^48^. Aβ may also indirectly lead to hyperconnectivity by promoting development of tau in the entorhinal cortex^24, 25^, which is associated with hyperactivity in medial temporal lobe^29, 49, 50^. Together, this previous literature in combination with our current results suggest that the hyperconnectivity between alEC and DG/CA3 observed in the present study may be a direct result of Alzheimer’s pathology, which preferentially disrupts alEC-DG/CA3 communication and expresses as deficits in object mnemonic discrimination.

We did not observe any statistically significant relationships between spatial mnemonic discrimination performance and entorhinal-hippocampal circuit FC. This is consistent with previous studies in cognitively normal older adults that failed to find relationships between spatial mnemonic discrimination and abnormal task-based activation in pmEC^16, 44^ or deficits in performance^16^ compared to young adults. However, a targeted analyses of the hypothesized connection between pmEC, which preferentially supports spatial processing^14^, and DG/CA3H revealed a nearly significant difference between high and low performing groups. It is likely that the connection between pmEC and DG/CA3, as well as performance on spatial mnemonic discrimination, is affected later in the development of AD.

Aβ pathology was also related to further hypoconnectivity and neurodegeneration within the hippocampus itself, particularly involving CA1 and Sub. Hypoconnectivity within the hippocampus may be a direct result of hyperconnectivity between entorhinal cortex and hippocampus. A study in mice found that Aβ-related excitatory increases in entorhinal cortex activation was associated with compensatory down-regulation of subiculum activity, which was proposed to be a protective response^51^. The connection between CA1 and Sub is of particular interest due to the propensity of tau pathology to target these subfields while sparing DG/CA3 until late stages of AD^18^. Future research should investigate specific tasks that probe CA1 and Sub in the context of aging and preclinical AD to determine if FC between these regions is associated with cognitive performance.

We did not find contributions of anterior-temporal and posterior-medial cortical networks to object or spatial mnemonic discrimination performance, respectively. Previous work has demonstrated a loss of functional specificity within these networks during object and scene memory in aging^12, 29^. Due to our high-resolution sequence with partial field of view focused on medial temporal lobe, we were unable to sample certain cortical regions commonly considered parts of these networks^32^. Disrupted FC in these networks may also arise later in disease progression or contribute more broadly to memory than specific deficits in mnemonic discrimination. Nonetheless, Aβ was associated with FC within the posteromedial network, which is consistent with previous findings^29, 41^.

There was no direct association between mnemonic discrimination performance and neurodegeneration. This suggests that FC, which can reflect both coordinated activation and structural connections, may be more sensitive to subtle cognitive changes. Previous studies have identified relationships between alEC volume/thickness and cognitive performance or status in older adults^45, 52, 53^. However, because changes in neural function occur before overt neurodegeneration, it is likely our results are detecting dysfunction that may not yet be observable with coarse measurements of structure such as volume.

Other mechanisms of age-related cognitive decline besides the development of Alzheimer’s pathology likely also play a role in impaired mnemonic discrimination in aging. In an outbred rat model that does not manifest AD pathological hallmarks, older rats show rigidity when exploring novel environments^6^, a change indicating a shift from pattern separation to pattern completion. This shift may result from functional and structural changes to DG and CA3 which increase engagement of autoassociative networks in CA3, biasing the computational balance towards pattern completion^4^. These changes have also been linked to circuit level changes such as decreased inhibition in the DG hilar region^54^ as well as loss of reelin protein and decreased synaptophysin immunoreactivity^55^. Future research should assess hippocampal function in older adults in the absence of Alzheimer’s proteinopathies to elucidate other mechanisms that may contribute to age-related memory decline.

This study has several limitations. First, we did not have tau-PET to evaluate tau within medial temporal lobe. While Aβ+ participants are highly likely to have entorhinal tau pathology^30^, some participants classified as Aβ- may also have age-related development of tau in medial temporal lobe known as primary age-related tauopathy^56^. Future studies should assess the role of tau on this circuit as high levels of tau may eventually lead to reduced rather than increased FC^48, 57^. Second, different ROIs were used to calculate volume, which may have diminished the strength of associations due the decreased specificity of the regions. Nonetheless, we observed associations between volume with both FC and Aβ pathology that were consistent with our hypotheses. Finally, 7T fMRI data could enable an even more precise measurement of distinct FC within subfield ROIs^5^. However, we applied semipartial correlations and no spatial smoothing to control for potential blurring of signal from adjacent ROIs, and found distinct patterns of FC that matched our hypotheses and previous literature. Replication of our findings at higher resolution may uncover additional novel associations.

In conclusion, our results suggest that Aβ pathology indirectly leads to mnemonic discrimination impairment through entorhinal-hippocampal circuit dysfunction. The vulnerability of the alEC to Alzheimer’s pathology may result in altered communication with DG/CA3, leading to impairment of object mnemonic discrimination early in the pathogenesis of AD. Aβ pathology further contributes to altered FC and neurodegeneration within the hippocampus itself, indicating more general dysfunction. Because resting state FC is easier to acquire than task-based fMRI and can be standardized across sites in large-scale studies, disruptions in FC within the entorhinal-hippocampal circuit could be a valuable biomarker of emerging memory decline in clinical trials.

## METHODS

### Participants

Cognitively normal older adults aged ≥60 years from the Biomarker Exploration in Aging, Cognition, and Neurodegeneration (BEACoN: NIA R01AG053555, PI: Yassa) study with high resolution resting state fMRI (N=89) were selected for the current study. After exclusion for fMRI data quality (i.e. signal drop out, field of view, motion; see below for details), data from 64 participants was analyzed. Inclusion criteria for BEACoN includes age ≥60 years, performance on cognitive assessments within age-adjusted normal range (within 1.5 standard deviations), no major health problems, co-morbid neurological disease, or significant psychiatric disorders, no use of medication for anxiety or depression or illicit drugs, and no MRI or PET contraindications. All participants provided written informed consent in accordance with the Institutional Review Board of the University of California, Irvine.

### MRI Acquisition

All participants received structural and resting state functional MRI at UCI on a 3T Prisma scanner (Siemens Medical System, Munich, Germany) equipped with a 32- channel head coil. A whole brain, high resolution T1-weighetd volumetric magnetization prepared rapid gradient echo images (MPRAGE) was acquired for structural analyses (voxel size = 0.8mm^3^ resolution, TR/TE/TI = 2300/2.38/902 ms, flip angle=8°, 240 slices acquired sagittally). High resolution T2*-weighted echo planar images (EPI) were acquired to assess functional connectivity (voxel size = 1.8mm^3^, TR/TE = 2500/26 ms, flip angle = 70°, 39 slices, R>>L phase encode, partial acquisition covering temporal lobe, 84 volumes). During acquisition, participants were instructed to remain awake and focus on a fixation cross on the screen. High resolution 3D T2-weighted turbo spin echo (TSE) images were acquired in oblique coronal orientation (voxel size = 0.4 x 0.4 mm in-plane resolution, slice thickness = 2mm, TR/TE = 5000/84 ms, 23 slices) for hippocampal segmentation and volumetry with ASHS.

### Structural MRI Processing

Structural T1 images were used for coregistration of both MRI and PET, and for segmentation of medial temporal subregions. T1 images were processed with Statistical Parametric Mapping (SPM, version 12, Wellcome Trust Center) and segmented into gray, white, and CSF compartments. T1 images were then skull stripped and warped to a study specific template using Advanced Normalization Tools (ANTs)^58^. T1 images were also processed with FreeSurfer v.6.0^59^ to obtain a native space regions of interest for florbetapir quantification.

To obtain measures of medial temporal lobe subregional volumes, both T1 and T2 structural images were processed using Automated Segmentation of Hippocampal Subfield (ASHS) software^60^. Segmentations were quality checked visually by trained researchers, and all passed inspection. Volumes of the entorhinal cortex, DG, CA3, CA1, and subiculum were used for analyses, and normalized by total intracranial volume (TIV) estimates derived from ANTs.

### Resting State Functional MRI Processing

Resting state fMRI data was preprocessed with SPM12 using a standard pipeline including slice time correction, realignment, and coregistration to the T1 structural image. No spatial smoothing was performed to maintain the high resolution of the images and enable more accurate quantification of signal within nearby subregions. Functional images were warped to a study-specific template with ANTs, applying the transformation parameters derived from structural warping.

#### ROIs

ROIs were defined using an in-house atlas of the temporal lobe and related regions, described in detail previously^13, 16, 31^. Briefly, the segmentation of hippocampal subfields^60, 61^, entorhinal subregions^14, 62^ (see Reagh et al., 2018), and cortical temporal regions^63^ was performed in accordance with standardized reliable protocols. The analysis included the following regions: anterolateral entorhinal cortex (alEC), posteromedial entorhinal cortex (pmEC), perirhinal cortex (PRC), parahippocampal cortex (PHC), amygdala (amyg), retrosplenial cortex (RSC), head and tail of dentate gyrus/CA3 (DG/CA3H, DG/CA3T), CA1 (CA1H, CA1T), and subiculum (SubH, SubT), bilaterally. ROIs across 10 coronal slices spanning the length of the ROI atlas are depicted in **Figure 2A**.

To ensure all ROIs had reliable signal due to the possibility of signal drop out in medial temporal lobe, we performed strict thresholding and exclusion of low signal. For each participant, mean signal across the gray matter was quantified, and voxels with <25% of the mean gray matter signal were removed. Participants with <50% of any ROI remaining were excluded from subsequent analyses due to lack of data, resulting in the exclusion of 13 participants. Two additional participants were excluded due to the partial acquisition field of view not including the full extent of the medial temporal lobe.

#### Denoising

Resting state fMRI data were optimized for functional connectivity analyses using the CONN toolbox^33^ (version 20) implemented in Matlab version 2019b (The MathWorks, Inc, Natick, MA). Outlier volumes were detected using Artifact Detection Tools (ART) implemented within CONN using conservative threshold of motion >0.5mm/TR and a global intensity z-score of 3. Ten subjects were flagged for >20% volumes detected as outliers and were removed from further analyses^29, 49^. Denoising was then performed, including six realignment parameters and their first-order derivatives (translations and rotations), spike regressors generated from outlier detection^64, 65^, anatomical CompCor^66^ (first five components of time series signal from white matter and CSF), bandpass filter [0.008-0.1 Hz], and linear detrending applied to the residual time series.

#### Functional Connectivity Analysis

Using CONN, denoised time series were extracted from each ROI, and semi-partial correlations were used to perform ROI-to-ROI functional connectivity analyses. Semi-partial correlations were chosen to control for any signal bleed in between spatially adjacent ROIs^34, 35^. Functional connectivity strength (Fisher’s r-to-z’ transformed correlation coefficient) was extracted for each ROI pair. Because we did not have specific hypotheses about hemispheric lateralization, and to reduce the number of statistical comparisons, we averaged functional connectivity strength between left and right ROI pairs for all analyses.

Second-level analyses were conducted in CONN. To determine patterns of group-level functional connectivity, we performed a between-subjects contrast using a one-sample t-test (two-tailed) for all seeds within each hemisphere (**Figure 2B**). Significant connections were determined with a connection-level threshold of p<0.05 p-uncorrected (two-sided) and a cluster level threshold of p<0.05 p-FDR corrected.

### Aβ PET

To quantify Aβ pathology, participants received ^18^F-florbetapir (FBP) PET performed on an ECAT High Resolution Research Tomograph (HRRT, CTI/Siemens, Knoxville, TN, USA) at CCNI. Ten mCi of tracer was injected, and four five-minute frames were collected between 50-70 min post-injection. FBP data was reconstructed with attenuation correction, scatter correction, and 2mm^3^ Gaussian smoothing. FBP images were then realigned, co-registered to the T1 MRI, and normalized by a whole cerebellum reference region to produce SUVR images. Additional 6mm^3^ Gaussian smoothing was then applied to achieve an effective resolution of 8mm^3^. The mean SUVR of a previously validated cortical composite region was quantified as a measure of global Aβ, and Aβ+ status was determined using a threshold of >1.11 SUVR^39^.

### Cognitive Data

#### Mnemonic Discrimination Task

Participants performed a mnemonic discrimination task (MDT) with both object (MDTO) and spatial (MDTS) versions (**Figure 1**), similar to previously described versions^16^. Briefly, each task consisted of a study phase, where participants are asked to make “Indoor/Outdoor” judgements about each stimulus (presented for 2 sec, ISI = 0.5 sec). In the object version, each object is presented at the center of the screen, while in the spatial version, each object is presented in a random grid position within the screen. After the study phase, participants complete a test phase. In the object version, the test phase consists of identical repeats of object stimuli (targets; correct response “old”) and lure stimuli (correct response “new”) which have either low or high visual similarity to the original object. In the spatial version, participants are tested on the spatial position of each object, with targets in the exact same location (correct response “same”), and lure stimuli (correct response “different”) with either low similarity (presented within a different quadrant of the grid), or high similarity (presented within a different position within the same quadrant) to the original target location. Participants were allowed 2 seconds to make a response before the next stimulus appeared. Each participant saw a unique order of stimuli for each phase.

As with all of our prior work using the MDT tasks, we calculated the response-bias corrected lure discrimination index (LDI) quantified as p(“New or Different” | Lure) – p (“New or Different” | Target) for the object and spatial task versions separately. We also calculated LDI specifically for highly similar object and spatial lures. We focused our analyses on the highly similar lure trials to tax pattern separation performance but also replicated analyses with all trial types combined. We classified participants into low and high performing groups based on a median split of high similarity LDI score on the object (median=0.185; low: MDTO-; high: MDTO+) and spatial (median=0.21; low: MDTS-; high: MDTS+) trials, separately.

#### Neuropsychological Assessment

Participants also performed standard neuropsychological assessment, including the Rey Auditory Verbal Learning Test (RAVLT) and the Mini Mental State Examination. The delayed recall measure of the RAVLT (A7) was used for control analyses testing general episodic memory, and participants were similarly grouped into low and high performers with a median split (median=11; low: RAVLT-; high: RAVLT+).

### Statistical Analyses

All statistical analyses were performed using jamovi v1.6 (https://www.jamovi.org) and RStudio v1.4. Functional connectivity differences across groups were assessed with repeated measures ANCOVAs, with functional connectivity (FC between ROI pairs) as a within-subjects factor, group based on performance or Aβ status (e.g. MDTO- vs. MDTO+, Aβ+ vs. Aβ-) as a between subjects factor, and age and sex as covariates of no interest. Main effects and interactions were considered significant at p<0.05 (two-tailed). We tested for main effects of performance/Aβ status to determine if functional connectivity was overall different between groups, and significant region by performance/Aβ status interactions to determine if groups varied in FC between specific ROI pairs. In cases of significant group by FC interactions, we further examined which ROI pairs were driving this interaction with post-hoc pairwise comparisons, considered significant at p<0.05 (two-tailed).

Bivariate and partial (controlling for age and sex) correlations were performed to assess relationships between continuous variables. Linear regressions were performed to test interactions between MDTO performance and neurodegeneration in predicting FC, including age and sex in the model. Independent samples t-tests (two-tailed) were also performed to assess group differences in neurodegeneration.

## Acknowledgments

This research was supported by NIA grants R01AG053555 (to M.A.Y.) and F32AG074621 (to J.N.A.) and the Alzheimer’s Disease Research Center at UC Irvine (P50 AG016573).

## Author Contributions

Conceptualization, JNA & MAY; Data Acquisition, ALH, AM, LM; Methodology & Software, JNA, SK, BR, MS, LT, DBK; Formal Analysis, JNA; Writing – Original Draft, JNA & MAY; Writing – Reviewing and Editing, all authors; Funding Acquisition, MAY.

## Data Availability

The data that support the findings of this study are available from the corresponding author upon reasonable request.

## REFERENCES

1. Leal, S. L. & Yassa, M. A. Integrating new findings and examining clinical applications of pattern separation. Nat. Neurosci. 21, 163–173 (2018).

2. Yassa, M. A. & Stark, C. E. L. Pattern separation in the hippocampus. Trends Neurosci. 34, 515–525 (2011).

3. Witter, M. P. The perforant path: projections from the entorhinal cortex to the dentate gyrus. Prog. Brain Res. 163, 43–61 (2007).

4. Wilson, I. A., Gallagher, M., Eichenbaum, H. & Tanila, H. Neurocognitive aging: prior memories hinder new hippocampal encoding. Trends Neurosci. 29, 662–670 (2006).

5. Berron, D. et al. Strong Evidence for Pattern Separation in Human Dentate Gyrus. J. Neurosci. 36, 7569–7579 (2016).

6. Wilson, I. A., Ikonen, S., Gallagher, M., Eichenbaum, H. & Tanila, H. Age-associated alterations of hippocampal place cells are subregion specific. J. Neurosci. 25, 6877–6886 (2005).

7. Yassa, M. A. et al. Pattern separation deficits associated with increased hippocampal CA3 and dentate gyrus activity in nondemented older adults. Hippocampus 21, 968–979 (2011).

8. Yassa, M. A. et al. High-resolution structural and functional MRI of hippocampal CA3 and dentate gyrus in patients with amnestic Mild Cognitive Impairment. Neuroimage 51, 1242–1252 (2010).

9. Bakker, A. et al. Reduction of Hippocampal Hyperactivity Improves Cognition in Amnestic Mild Cognitive Impairment. Neuron 74, 467–474 (2012).

10. Reagh, Z. M. et al. Greater loss of object than spatial mnemonic discrimination in aged adults. Hippocampus 26, 417–422 (2016).

11. Güsten, J., Ziegler, G., Düzel, E. & Berron, D. Age impairs mnemonic discrimination of objects more than scenes: A web-based, large-scale approach across the lifespan. Cortex 137, 138–148 (2021).

12. Berron, D. et al. Age-related functional changes in domain-specific medial temporal lobe pathways. Neurobiol. Aging 65, 86–97 (2018).

13. Reagh, Z. M. & Yassa, M. A. Object and spatial mnemonic interference differentially engage lateral and medial entorhinal cortex in humans. Proc. Natl. Acad. Sci. U. S. A. 111, E4264–73 (2014).

14. Maass, A., Berron, D., Libby, L., Ranganath, C. & Düzel, E. Functional subregions of the human entorhinal cortex. Elife 4, 1–20 (2015).

15. Knierim, J. J., Neunuebel, J. P. & Deshmukh, S. S. Functional correlates of the lateral and medial entorhinal cortex: Objects, path integration and local - Global reference frames. Philos. Trans. R. Soc. B Biol. Sci. 369, (2014).

16. Reagh, Z. M. et al. Functional imbalance of anterolateral entorhinal cortex and hippocampal dentate/CA3 underlies age-related object pattern separation deficits. Neuron 97, 1187–1198.e4 (2018).

17. Braak, H. & Braak, E. On areas of transition between entorhinal allocortex and temporal isocortex in the human brain. Normal morphology and lamina-specific pathology in Alzheimer’s disease. Acta Neuropathol. 68, 325–332 (1985).

18. Braak, H. & Braak, E. Neuropathological stageing of Alzheimer-related changes. Acta Neuropathol 82, 239–259 (1991).

19. Palop, J. J. & Mucke, L. Amyloid-beta-induced neuronal dysfunction in Alzheimer’s disease: from synapses toward neural networks. Nat. Neurosci. 13, 812–818 (2010).

20. Busche, M. A. et al. Clusters of hyperactive neurons near amyloid plaques in a mouse model of Alzheimer’s Disease. Science (80-.). 321, 1686–1689 (2008).

21. Huijbers, W. et al. Amyloid-β deposition in mild cognitive impairment is associated with increased hippocampal activity, atrophy and clinical progression. Brain 138, 1023–1035 (2015).

22. Mormino, E. C. et al. Aβ Deposition in aging is associated with increases in brain activation during successful memory encoding. Cereb. Cortex 22, 1813–1823 (2012).

23. Huijbers, W. et al. Amyloid deposition is linked to aberrant entorhinal activity among cognitively normal older adults. J. Neurosci. 34, 5200–5210 (2014).

24. Adams, J. N., Harrison, T. M., Maass, A., Baker, S. L. & Jagust, W. J. Distinct Factors Drive the Spatiotemporal Progression of Tau Pathology in Older Adults. J. Neurosci. 42, 1352–1361 (2022).

25. Sanchez, J. S., et al. The cortical origin and initial spread of medial temporal tauopathy in Alzheimer ’ s disease assessed with positron emission tomography. 0655, (2021).

26. Jack, C. R. et al. Hypothetical model of dynamic biomarkers of the Alzheimer’s pathological cascade. Lancet Neurol. 9, 119–128 (2010).

27. Berron, D. et al. Higher CSF tau levels are related to hippocampal hyperactivity and object mnemonic discrimination in older adults. J. Neurosci. 39, 1279–19 (2019).

28. Marks, S. M., Lockhart, S. N., Baker, S. L. & Jagust, W. J. Tau and β-amyloid are associated with medial temporal lobe structure, function and memory encoding in normal aging. J. Neurosci. 37, 3769–16 (2017).

29. Maass, A. et al. Alzheimer’s pathology targets distinct memory networks in the ageing brain. Brain 142, (2019).

30. Braak, H. & Braak, E. Frequency of stages of Alzheimer-related lesions in different age categories. Neurobiol. Aging 18, 351–357 (1997).

31. Yassa, M. A., Mattfeld, A. T., Stark, S. M. & Stark, C. E. L. Age-related memory deficits linked to circuit-specific disruptions in the hippocampus. Proc. Natl. Acad. Sci. U. S. A. 108, 8873–8878 (2011).

32. Ranganath, C. & Ritchey, M. Two cortical systems for memory-guided behaviour. Nat. Rev. Neurosci. 13, 713–726 (2012).

33. Whitfield-Gabrieli, S. & Nieto-Castanon, A. Conn: a functional connectivity toolbox for correlated and anticorrelated brain networks. Brain Connect. 2, 125–141 (2012).

34. Adams, J. N., Maass, A., Harrison, T. M., Baker, S. L. & Jagust, W. J. Cortical tau deposition follows patterns of entorhinal functional connectivity in aging. Elife 8, 1–22 (2019).

35. Dalton, M. A., McCormick, C., De Luca, F., Clark, I. A. & Maguire, E. A. Functional connectivity along the anterior–posterior axis of hippocampal subfields in the ageing human brain. Hippocampus 1049–1062 (2019) doi:10.1002/hipo.23097.

36. Amaral, D. G. & Lavenex, P. Hippocampal Neuroanatomy. in The Hippocampus Book (eds. Andersen, P., Morris, R., Amaral, D., Bliss, T. & O’Keefe, J.) vol. 15 (Oxford University Press, 2006).

37. Gordon, B. A., Blazey, T., Benzinger, T. L. S. & Head, D. Effects of Aging and Alzheimer’s Disease Along the Longitudinal Axis of the Hippocampus. J Alzheimers Dis. 37, (2013).

38. Das, S. R. et al. Increased functional connectivity within medial temporal lobe in mild cognitive impairment. Hippocampus 23, 1–6 (2013).

39. Landau, S. M. et al. Amyloid deposition, hypometabolism, and longitudinal cognitive decline. Ann. Neurol. 72, 578–586 (2012).

40. Berron, D., van Westen, D., Ossenkoppele, R., Strandberg, O. & Hansson, O. Medial temporal lobe connectivity and its associations with cognition in early Alzheimer’s disease. Brain (2020) doi:10.1093/brain/awaa068.

41. Cassady, K. E. et al. Alzheimer’s Pathology Is Associated with Dedifferentiation of Intrinsic Functional Memory Networks in Aging. Cereb. Cortex 1–13 (2021) doi:10.1093/cercor/bhab122.

42. Bennett, I. J. & Stark, C. E. L. Mnemonic discrimination relates to perforant path integrity: An ultra-high resolution diffusion tensor imaging study. Neurobiol. Learn. Mem. 129, 107–112 (2016).

43. Burke, S. N., Turner, S. M., Desrosiers, C. L., Johnson, S. A. & Maurer, A. P. Perforant path fiber loss results in mnemonic discrimination task deficits in young rats. Front. Syst. Neurosci. 12, 1–13 (2018).

44. Berron, D. Higher CSF tau levels are related to hippocampal hyperactivity and object mnemonic discrimination in older adults. J. Neurosci. (2019).

45. Tran, T. T., Speck, C. L., Gallagher, M. & Bakker, A. Lateral Entorhinal Cortex Dysfunction in Amnestic Mild Cognitive Impairment. Neurobiol. Aging (2021) doi:10.1016/j.neurobiolaging.2021.12.008.

46. Sinha, N. et al. ABCA7 Risk Variant in Healthy Older African Americans is Associated with a Functionally Isolated Entorhinal Cortex Mediating Deficient Generalization of Prior Discrimination Training. Hippocampus 29, 527–538 (2019).

47. Dautricourt, S. et al. Longitudinal Changes in Hippocampal Network Connectivity in Alzheimer’s Disease. Ann. Neurol. 90, 391–406 (2021).

48. Schultz, A. P. et al. Phases of Hyperconnectivity and Hypoconnectivity in the Default Mode and Salience Networks Track with Amyloid and Tau in Clinically Normal Individuals. J. Neurosci. 37, 4323–4331 (2017).

49. Adams, J. N. et al. Reduced repetition suppression in aging is driven by tau-related hyperactivity in medial temporal lobe. J. Neurosci. 41, JN-RM-2504-20 (2021).

50. Huijbers, X. W. et al. Tau accumulation in clinically normal older adults is associated with hippocampal hyperactivity. J. Neurosci. 39, 548–556 (2019).

51. Angulo, S. L. et al. Tau and amyloid-related pathologies in the entorhinal cortex have divergent effects in the hippocampal circuit. Neurobiol. Dis. 108, 261–276 (2017).

52. Olsen, R. K. et al. Human anterolateral entorhinal cortex volumes are associated with cognitive decline in aging prior to clinical diagnosis. Neurobiol. Aging 57, 195–205 (2017).

53. Holbrook, A., Tustison, N., Marquez, F., Roberts, J. & Michael, A. Anterolateral entorhinal cortex thickness as a new biomarker for early detection of Alzheimer ’ s disease Abstract : 1–27 (2019).

54. Spiegel, A. M., Koh, M. T., Vogt, N. M., Rapp, P. R. & Gallagher, M. Hilar interneuron vulnerability distinguishes aged rats with memory impairment. J. Comp. Neurol. 521, 3508–3523 (2013).

55. Stranahan, A. M., Haberman, R. P. & Gallagher, M. Cognitive decline is associated with reduced reelin expression in the entorhinal cortex of aged rats. Cereb. Cortex 21, 392–400 (2011).

56. Crary, J. F. et al. Primary age-related tauopathy (PART): a common pathology associated with human aging. Acta Neuropathol. 128, 755–766 (2014).

57. Busche, M. A. et al. Tau impairs neural circuits, dominating amyloid-β effects, in Alzheimer models in vivo. Nat. Neurosci. 22, 57–64 (2019).

58. Tustison, N. J. et al. The ANTsX ecosystem for quantitative biological and medical imaging. Sci. Rep. 11, 9068 (2021).

59. Fischl, B. FreeSurfer. Neuroimage 62, 774–781 (2012).

60. Yushkevich, P. A. et al. Automated volumetry and regional thickness analysis of hippocampal subfields and medial temporal cortical structures in mild cognitive impairment. Hum. Brain Mapp. 36, 258–287 (2015).

61. Wisse, L. E. M. et al. A harmonized segmentation protocol for hippocampal and parahippocampal subregions: why do we need one and what are the key goals? Hippocampus 27, 3–11 (2017).

62. Navarro Schröder, T., Haak, K. V, Zaragoza Jimenez, N. I., Beckmann, C. F. & Doeller, C. F. Functional topography of the human entorhinal cortex. Elife 4, 1–17 (2015).

63. Insausti, R. et al. MR volumetric analysis of the human entorhinal, perirhinal, and temporopolar cortices. Am. J. Neuroradiol. 19, 659–671 (1998).

64. Lemieux, L., Salek-Haddadi, A., Lund, T. E., Laufs, H. & Carmichael, D. Modelling large motion events in fMRI studies of patients with epilepsy. Magn. Reson. Imaging 25, 894–901 (2007).

65. Power, J. D., Schlaggar, B. L. & Petersen, S. E. Recent progress and outstanding issues in motion correction in resting state fMRI. Neuroimage 105, 536–551 (2015).

66. Behzadi, Y., Restom, K., Liau, J. & Liu, T. T. A component based noise correction method (CompCor) for BOLD and perfusion based fMRI. Neuroimage 37, 90–101 (2007).

